# From contigs towards chromosomes: automatic Improvement of Long Read Assemblies (ILRA)

**DOI:** 10.1101/2021.07.30.454413

**Authors:** José L. Ruiz, Susanne Reimering, Juan David Escobar-Prieto, Nicolas M. B. Brancucci, Diego F. Echeverry, Abdirahman I. Abdi, Matthias Marti, Elena Gómez-Díaz, Thomas D. Otto

## Abstract

Recent advances in long read technologies not only enable large consortia to aim to sequence all eukaryotes on Earth, but they also allow individual laboratories to sequence their species of interest with relatively low investment. Although there is a promise of long read technologies to obtain “perfect genomes”, the number of contigs often exceeds the number of chromosomes by far, containing many insertion and deletion errors around homopolymer tracks. To overcome these issues, we implemented the ILRA pipeline to correct long read-based assemblies, so contigs are reordered, renamed, merged, circularized, or filtered if erroneous or contaminated, and Illumina reads are used to correct homopolymer errors. We successfully tested our approach by improving the genomes of *Trypanosoma brucei* and *Leptosphaeria* spp, and generated four novel *Plasmodium falciparum* assemblies from field samples. We found that correcting homopolymer tracks reduced the number of genes incorrectly annotated as pseudogenes, but an iterative correction seems to be required to correct larger numbers of sequencing errors. In summary, we described and compared the performance of our new tool, which improved the quality of novel long read assemblies of genomes up to 1Gbp.

**Availability:** The tool is available at GitHub: https://github.com/ThomasDOtto/ILRA.

## INTRODUCTION

Challenges of genome assemblies include sample quality, sequencing depth, number of repeats in the genomes and low complexity regions, which result in higher number of contigs or low consensus quality. Next Generation Sequencing techniques have undergone impressive development over the last few years, achieving unparalleled resolution and performance (Marx, 2021). If enough high molecular weight DNA and funding is available, the long read technologies Pacific Bioscience (PacBio) (Eid, et al., 2009) and Oxford Nanopore Technologies (ONT) (Branton, et al., 2008), combined with scaffolding methods such as HiC and Bionano, produce continuous sequences, as complex repeats can be spanned. Complementarily, short reads can be used to address limitations in long read sequencing, mainly by correcting homopolymer tracks and di/tri-mer repeats. These advances, together with the drop of price per base, have motivated the formation of large consortia, such as the Earth BioGenome Project that aims to sequence all eukaryotes on Earth (Lewin, et al., 2018). Beyond larger genome sizes, these consortia also include organisms of medium (up to 1 Gbp) and small (∼50 Mbp) genomes. To generate telomere-to-telomere assemblies with less than 1 error per 10,000 bases as gold standard reference genomes (Chain, et al., 2009; Koepfli, et al., 2015), the consortia use large quantities of high molecular weight and high-quality DNA, together with a range of sequencing technologies (short and long reads) and scaffolding methods. On the other hand, individual research groups tend to apply whole genome sequencing to either produce *de novo* assemblies of a few species of interest or to evaluate genome variation (i.e. single nucleotide and copy number variants, SNV-CNVs) within a species. As their resources are typically limited, in general the obtained assemblies are more fragmented, and more automatic finishing is needed.

These finishing steps have traditionally taken as much time and manual effort as prior sequencing and assembly, but since the advent of Illumina sequencing, several steps can now be automated with tools like PAGIT (Swain, et al., 2012). For long read assemblies, improving the sequence quality (i.e. polishing) is particularly needed, as the ONT and PacBio technologies cannot capture correctly repeats of 1-3 base pairs (homopolymer tracks and di/tri-mer repeats) (Koren, et al., 2019; Watson and Warr, 2019). Recent breakthroughs including the up-to-date PacBio Hi-Fi, ONT ultralong reads, Illumina Infinity or Ultima Genomics, do not necessarily generate telomere-to-telomere chromosomes, nor address errors around homopolymer tracks (Booeshaghi and Pachter, 2022). This issue is more pronounced for sequences with skewed base compositional bias. For example some pathogens, such as the malaria parasite *Plasmodium falciparum* with a GC content of 19%, resulting in large low complexity regions. Errors around homopolymer tracks affects the accuracy of the predicted gene models in all organisms. Therefore, tools such as iCORN2 (Otto, et al., 2010) and Pilon (Walker, et al., 2014) have been specifically developed to map Illumina short reads to correct genome assemblies, small errors and frameshifts. While various assembler software exist, pipelines to streamline finishing tools are still limited (Swain, et al., 2012). Assemblosis (Korhonen, et al., 2019), ARAMIS (Sacristan-Horcajada, et al., 2021) and MpGAP (Marques de Almeida and Pappas, 2022) are recent examples of the very few pipelines available to automatically assemble long reads and polish the resulting sequences. Their limitations include the use of specific software, such as particular assembler or error correction tools, and they may not be easy to install and run. Furthermore, these pipelines could be improved by adding more steps such as resolving overlapping contigs, excluding noise, using naming conventions, or expediting the process of uploading to external databases.

In this study, we overcome the limitations above and further streamline the automatic finishing process, implementing a more integrative approach rather than individual polishing steps. We developed the automatic Improvement of Long Read Assemblies (ILRA), an easy-to-use pipeline that combines novel and existing tools to clean *de novo* genome assemblies. We have tested the IRLA pipeline on human data to understand the limitation of genome size and then applied it to several genomes with varying sequencing depth, median read length and sequencing approaches, including four novel *Plasmodium falciparum* genomes, a *Trypanosoma brucei* assembly by PacBio (Muller, et al., 2018), and two fungi assemblies (*Leptosphaeria* spp.) by ONT (Branton, et al., 2008). We conclude that ILRA is generally applicable to long read assemblies across species and outperforms existing tools. Overall, we provide a valuable resource that could be widely implemented by non-specialists in many ongoing and future sequencing projects, especially those focusing on smaller and/or challenging genomes.

## METHODS

### Assemblies and annotation

To determine whether assemblers are generating sequences with further need of improvements, different assembler software were compared: HGAP (Chin, et al., 2013), Canu (Koren, et al., 2017) or Wtdbg2 (Ruan and Li, 2020), which are all using PacBio long reads, and MaSuRCA (Zimin, et al., 2013), which combines both PacBio long reads and Illumina short reads. As input, we used four sets of reads for three different *P. falciparum* isolates (PfCO01, PfKE07 and Pf2004; Table 1 and Supplementary Table 1). Specific challenges included: DNA for the PfKE07 sample was amplified by whole genome amplification (WGA), and the same library for the PfCO01 sample was sequenced using two different PacBio sequencers, the RSII and the Sequel.

**Table 1:** Overview of *P. falciparum* assemblies by MaSuRCA and information of the datasets used in this study. Standard - normal DNA preparation.

| Isolates | Pf3D7<br>reference | PfCO01 | PfCO01 | PfKE07 | Pf2004 |
| --- | --- | --- | --- | --- | --- |
| **Library preparation** | - | Standard | Standard | Whole Genome<br>Amplification | Standard |
| **Sequencer machine**<br>**(PacBio)** | - | RSII | Sequel | RSII | RSII |
| **Number of contigs** | 16 | 30 | 41 | 259 | 21 |
| **Size (Mb)** | 23.33 | 23.36 | 23.43 | 21.76 | 23.30 |
| **Number of genes** | 5,720 | 6,145 | 6,210 | 5,067 | 5,701 |
| **Number of pseudogenes** | 158 | 154 | 163 | 449 | 156 |

HGAP was run with its default parameters, setting the genome size to 23.5 Mbp. We used version 3 for all assemblies, except for PfCO01 where version 4 was used for the Sequel reads. Canu v2.2 was run with parameters -pacbio-raw, corMaxEvidenceErate=0.15, genomeSize=24m. Wtdbg2 v2.5 was run with -g 24m, -x rs for PfKE07 and the run of PfCO01 from RSII, and -x sq for the alternative run of PfCO01 from Sequel. MaSuRCA v4.1.0 was run with default parameters, and Illumina short reads (specifying mean=700 and stdev=100) were also provided. The assemblies by Wtdbg2 were preliminary polished using the Illumina short reads, following the recommendations by the authors in the official GitHub repository. All jobs were run on the same machine, with 60 cores and 250 GB of RAM available.

All the assemblies in this study were annotated using the Companion (Steinbiss, et al., 2016) web server version 2022. We used *P. falciparum* 3D7 and *T. brucei* TREU927 as references for the corresponding assemblies. For the fungal assemblies (*Leptosphaeria* spp.), we first used the interactive Tree of Life (Letunic and Bork, 2019) and identified amongst the available references in Companion *Fusarium verticillioides* as the closest fungal species. Hence, *F. verticillioides* was used as reference for annotation. Companion default parameters were used in all cases, except for AUGUSTUS score threshold = 0.2, and taxon ID = 5833, 5691, 5022 and 220672 for *P. falciparum, T. brucei, L. maculans* and *L. biglobosa*, respectively. Pseudochromosome contiguation was deactivated for the fungal assemblies for which Companion only provided close references. When available, the numbers of annotated genes and pseudogenes in the *P. falciparum* and *T. brucei* reference genomes were consulted in GeneDB (Logan-Klumpler, et al., 2012), within the release 58 of PlasmoDB and TriTrypDB (Amos, et al., 2022). Full statistics and information for all the assemblies in this study are shown in Supplementary Table 1.

Visualizations and analyses comparing the assemblies to each other and to the references were performed using Artemis and the Artemis Comparison Tool (ACT) v16.0.9 (Carver, et al., 2008). Primary assembly statistics were obtained using the software assembly-stats (https://github.com/sanger-pathogens/assembly-stats).

### Comparison of iCORN2 and Pilon correction

iCORN2 v1.0 and Pilon v1.24 were evaluated for their performance and accuracy of corrections of small indels and one base pair errors in long read assemblies. As a truth set for the experiment, the uncorrected long read assembly of *P. falciparum* (Pf3D7, available at ftp://ftp.sanger.ac.uk/pub/project/pathogens/Plasmodium/falciparum/PF3K/ReferenceGenomes_Version1/Pf3D7_PacbioAssemblies/) was used by both tools and compared to the current Pf3D7 reference (version v3.1). Illumina short reads of different lengths, 75 bp to 300 bp, were evaluated. The tools were run with the following parameters: 500 bp fragment length and 5 iterations for iCORN2, and for Pilon we provided aligned reads and used default options. Two different approaches were also implemented for each tool:

1. Mapping Illumina short reads with the bwa-mem2 v2.2.1 (Li, 2013) aligner with default parameters for Pilon. iCORN2 used its default aligner SMALT v0.7.6 (https://github.com/rcallahan/smalt).
2. Mapping Illumina short reads with more sensitivity using Bowtie2 v2.4.5 (Langmead and Salzberg, 2012) with parameters -X 1200 --very-sensitive -N 1 -L 31 --rdg 5,2.

To evaluate the corrections, MEGAblast v2.2.26 (Morgulis, et al., 2008) with parameters -F F - e 1e-80 -m 8 was used to analyze syntenic hits of length ≥ 10 kb, focusing on a single region in chromosome 5 (Supplementary Table 2). The results for all chromosomes were processed manually to approximately identify the same regions in the different comparisons between the reference genome and the differentially corrected assemblies, and to evaluate the correction steps by showing final aggregated values (Table 2 and Supplementary Table 2). We used the Companion web server version 2022 to annotate the genomes and to assess the different number of pseudogenes after the independent correction steps by both tools.

**Table 2:** Comparison of Pilon with iCORN2. A published Pf3D7 PacBio assembly by HGAP was corrected with Pilon and with 5 iterations of iCORN2 (mapping of Illumina short reads by Bowtie2 for both tools). The results were compared against the *P. falciparum* 3D7 reference using MegaBLAST to obtain the number of differences between the reference and corrected sequence. The number of pseudogenes in the Pf3D7 reference is 158.

| Read length | Tool | Indels | SNPs | Pseudogenes |
| --- | --- | --- | --- | --- |
| **75bp** | Pilon | 26,037 | 1,973 | 531 |
|  | iCORN2 | 9,768 | 1,961 | 164 |
| **100bp** | Pilon | 17,905 | 1,879 | 255 |
|  | iCORN2 | 6,402 | 2,235 | 143 |
| **300bp** | Pilon | 19,473 | 1,749 | 197 |
|  | iCORN2 | 4,485 | 3,919 | 141 |

For the evaluation of the correction of a *T. brucei* genome, we used an uncorrected long read assembly (Muller, et al., 2018) and two sets of Illumina short reads that we concatenated (Supplementary Table 3). As truth set, we used the *T. brucei* Lister strain 427 2018 reference, using MEGAblast as described above. We kept the hits to the chromosomes in the reference genome larger than 10 kb and with identity > 99%. The results were processed manually to take unique alignments, prioritizing larger hits with higher bit-scores. The output for the comparison of the corrections of the *T. brucei* assembly is provided in Supplementary Table 3.

**Table 3:** Final comparison of Pilon vs. iCORN2 using the Bowtie2 aligner to map Illumina short reads, 5 iterations, and comparing the number of annotated pseudogenes.

| Organism | Tool<br>(5 iterations) | Pseudogenes | Runtime<br>(hours) |
| --- | --- | --- | --- |
| *P. falciparum* | Pilon | 210 | 1.91 |
|  | iCORN2 | 143 | 9.04 |
| *T. brucei* | Pilon | 5,037 | 10.51 |
|  | iCORN2 | 5,179 | 19.34 |

### External assembly and polishing pipelines

Assemblosis, ARAMIS and MpGAP were run on two of our novel *P. falciparum* datasets (Pf2004 and PfCO01 from RSII reads), and the output was compared to the results of ILRA (Supplementary Table 1). In the case of Assemblosis, assembly was performed by Canu and polishing was based on Pilon using short reads. Default parameters were used for Assemblosis, except for specifying 23m as the genomeSize, 3000 as the minReadLen and 0.15 as the corMaxEvidenceErate parameters. Following the guidelines by the authors of ARAMIS, the correction.sh script specifying the parameter -i 0.5 was applied to the alignment of the Illumina short reads to the assembly, obtained using bwa-mem2 (REF) with default parameters. Finally, MpGAP was used with default parameters except for 23.3m as genome_size and hybrid_strategy 2, so multiple assembly approaches for long reads were tested (i.e. Canu, Wtdbg2, Shasta, Raven, Flye or Unicycler) and then Pilon-based polishing was performed, implementing the Unicycler-polish program to iteratively correct until changes are determined to be minimal. We chose the best assemblies (i.e. Pilon-corrected assemblies from Flye) to integrate in this study.

### ILRA methods

We implemented ILRA as a bash script to automatically improve long reads assemblies. Supplementary Figure 1 shows the overview of the pipeline. The full information for the assemblies in this study and the statistics for the correction by ILRA are in Supplementary Table 1. The following steps are automatically performed by ILRA:

#### 1. Cleaning contigs

Contigs smaller than 5 kb are removed, as the mean read length for PacBio technologies is over 7 kb and small contigs are considered the product of inadequate reads, for example chimeric reads. Contained contigs that fully overlap with the sequence of larger contigs are also removed, since they may represent partially phased regions of the genome or mistakes by the assembler software. The overlap between contigs is determined using MegaBLAST with parameters -W 40 -F F -m 8 -e 1e-80. Finally, the contigs overlapping by more than 90% of their length and displaying more than 99% identity with any other contig are also removed. The names of all excluded contigs are stored in the file *Excluded*.*contigs*.*fofn*.

#### 2. Merging contigs

Overlapping contigs are merged if they have an overlapping region larger than 2 kb, they display identity larger than 99% and the coverage of the Illumina short reads is 40-60% of the median coverage in the full assembly. These requisites ensure that contigs are not merged due to repeats.

#### 3. Ordering contigs against a reference (optional)

The sequences are renamed and the contigs are reordered and orientated using a reference genome via ABACAS2 (Assefa, et al., 2009), specifying by default minimum alignment length 1 kb, 98% identity cut-off and ABA_CHECK_OVERLAP = 0. If no reference is provided, this step is omitted and the sequences get renamed later. Here, we used the Pf3D7 reference v3.1 (Bohme, et al., 2019) for the improvement of the four *P. falciparum* assemblies. To avoid ordering contigs in polymorphic regions, we just used the core region of the genome as reference (Otto, et al., 2018). We also improved a *T. brucei* assembly by PacBio using as reference the Lister strain 427 2018 (Muller, et al., 2018), and two fungi assemblies by ONT (*Leptosphaeria maculans* Nz-T4 and *L. biglobosa* G12-14) (Dutreux, et al., 2018) using previously published genomes as references (Grandaubert, et al., 2014). We extensively updated ABACAS2 to be included in ILRA, mainly by improving performance and multi-threading implementing GNU’s parallel. Whether to perform blasting and the number of contigs to process in parallel can be input by the user.

#### 4. Correcting homopolymer errors

ILRA corrects single-base discrepancies and indels, which are common shortcomings in long reads due to the presence of homopolymer tracks using Illumina short reads. ILRA uses by default iCORN2, with 500 bp fragment length and 3 iterations. Alternatively, users can choose to execute iterations of Pilon, with default parameters. In both cases, the user can specify the amount of RAM memory available so the programs would also run on machines with less resources available, at the expense of longer computation times. Here, we implemented a modified version of iCORN2 (v1.0). The main changes that we introduced are the use of Bowtie2 for read mapping to increase sensitivity (parameters -X 1200 --very-sensitive -N 1 -L 31 -- rdg 5,2), and the parallel processing of chunks of contigs (lower than ∼60 Mbp of size) to reduce execution time and memory usage. The number of contigs to be processed in parallel can be provided by the user depending on the available resources. This version has been made available in the main iCORN2 repository (http://icorn.sourceforge.net/). The tool requires Java runtime v1.7. The numbers of corrected SNPs and indels are also provided by iCORN2. In the case of Pilon, the execution within ILRA includes the more sensitive Bowtie2 command above, and the user can also provide the number of iterations to perform and the limit of memory usage, which is an approach not natively supported by Pilon. Additionally, the latest update of iCORN2 also allows for the use of Pilon for the processing of contigs or sequences larger than ∼60 Mbp of size, which previous versions of iCORN2 could not process.

#### 5. Circularizing plasmids

The genomes of organelles and extrachromosomal plasmids are circular, and the correct sequences need to be generated starting at the origin. Thus, the sequences corresponding to the mitochondria (or the apicoplast in some parasites such as *Plasmodium*, or the *T. brucei* “maxicircle” contig), are circularized by ILRA using Circlator v1.5.5 (Hunt, et al., 2015) with default parameters (command “all”). The corrected long reads provided by the assembler software and mapped to the assemblies are required by Circlator. ILRA provides the alignment using winnowmap(Jain, et al., 2022) v2.0.3 with default parameters (except for -x map-pb for PacBio reads and –x map-ont for ONT reads). ILRA also allows adaptation by the users, so it can be specified which sequences should be circularized.

#### 6. Decontaminating contigs

Contigs that are products of contaminations are identified by ILRA using a taxonomic classification approach by Kraken2 v2.1.2 (Wood, et al., 2019) with default parameters and a database using viral, plasmid, bacteria, archaea, fungi and UniVec sequences, together with multiple VEuPathDB databases, including PlasmoDB or TriTrypDB. Further sequence sets, such as the NCBI nucleotide non-redundant database or custom sequences, can be provided by the user to achieve better sensitivity, at the expense of larger space or memory usage and processing times. Decontamination is recommended prior to the *de novo* assembly process, but ILRA has been designed at this step to filter the contigs that are assigned to NCBI taxons representing potential contaminants. Taxonomic classification is assessed using the tools Recentrifuge v1.3.1 (Marti, 2019) with default parameters. To allow adaptation by the users, the NCBI taxon IDs to keep are provided to ILRA as an inline parameter so the organisms of interest are extracted by KrakenTools v1.2 (Wood and Salzberg, 2014).

ILRA can additionally perform BLAST of the final assembly against multiple databases, such as common contaminants, vector sequences, bacterial insertion sequences or ribosomal RNA genes. The aim is to follow the NCBI’s Foreign Contamination Screen: to mask some regions, together with renaming and reformatting some contigs if needed (*e*.*g*., adding the cell location), so the requirements of the most popular online databases (*i*.*e*. DDBJ/ENA/Genbank) are fully fulfilled. Thus, the ILRA-corrected assemblies may be directly uploaded to databases with minimal need of user input.

#### 7. Gathering the assembly statistics

Once the sequences have been processed and automatically polished, telomere-associated sequences and telomere repeats are counted to assess the completeness of the contigs and identify the potential chromosomes with both telomeres attached.

The sequences to analyze within the telomeres are set by default but can be also provided by the users as inline parameters. To evaluate the quality of the assemblies, ILRA also reports general statistics pre- and post-correction by multiple software inputs, including sequencing depth, read lengths, contig sizes, GC-content, assembly sizes, N50 values or number of gaps. Optionally, if a reference genome and an annotation file (GFF format) are provided, the software QUAST v5.2.0 (Mikheenko, et al., 2018) is also used to compute various metrics and plots, such as structural variants compared to the reference, misassemblies or mismatches. Here, for assessing the quality of the ILRA-corrected assemblies, we downloaded the annotation for *P. falciparum 3D7* and *T. brucei* Lister strain 427 2018 from GeneDB and the SiegelLab. ILRA also includes the tools plotsr v0.5.4 (Goel and Schneeberger, 2022), to plot synteny and structural rearrangements between the assembly and a reference genome, if provided, and BUSCO v5.4.0 (Simao, et al., 2015) to automatically assess the assembly completeness and to report various quality metrics.

### Sequence datasets

To develop and test our pipeline, we used three species with different sets of reads (Supplementary Table 1).

1. Through the Pf3K project reference (https://www.malariagen.net/projects/Pf3k), we generated several *Plasmodium falciparum* read sets using different library generation approaches, including WGA, Pacbio Sequel and PacBio RSII. More information on ethical and sequencing information are in the Supplementary Material.
2. *T. brucei* uncorrected PacBio long reads assembly (Muller, et al., 2018).
3. *L. maculans* Nz-T4 and *L. biglobosa* G12-14 ONT long reads assemblies (Dutreux, et al., 2018).

The fungi datasets were used because they were originally sequenced by ONT and the assemblies were polished twice with Pilon (Dutreux, et al., 2018). The uncorrected *T. brucei* genome was made available to us upon request. The *P. falciparum* reads sets covered a wider range of sequencing reads types and preparation methods, while representing a challenging genome in which we have expertise. As a proof of concept, we also tested the limits of ILRA by manually taking chunks of the human genome HG00733/GCA_003634875, which is fragmented in 566 contigs summing 5.8 Gbp (Garg, et al., 2021). From these sequences, we extracted subsets of different sizes and numbers of contigs: ∼100 Mbp, ∼150 Mbp, ∼300 Mbp, 500 Mbp and 1 Gbp, composed of 518, 521, 525, 528 and 535 contigs, respectively. We then corrected these test sequences with ILRA using the corresponding subsets of Illumina reads (SRR5535410) aligned to the chunks (Bowtie2 with default parameters except for -X 800). As a reference similar to the human genome, we provided a chimpanzee genome (GCF_002880755) removing unassigned scaffolds.

## RESULTS

In this study, we developed a new pipeline for the Improvement of Long Reads Assembly (ILRA). We first show that independently of the choice of assembler software, polishing seems to be always necessary. We then compared two tools to correct sequencing errors, Pilon and iCORN2, with the aim of including them in our pipeline to improve draft long read assemblies. Next, we applied the ILRA pipeline to seven assemblies of PacBio and ONT reads from the three studied species. Finally, we also compared our results with other assembly and polishing pipelines and explored possible limitations, such as genome sizes.

### Comparison of assembler software

First, we evaluated the impact of the use of different assemblers on the number of contigs and the presence of potential frameshifts due to homopolymer and di/tri-mer errors. We used PacBio reads from RSII and Sequel of different qualities (PfCO01 RSII, PfCO01 Sequel, Pf2004), and one dataset (PfKE07) whose RSII reads were generated from a library subjected to Whole Genome Amplification (WGA) (Supplementary Table 1). The four sets of *P. falciparum* reads were assembled with four different tools (HGAP, Canu, MaSuRCA and Wtdbg2) to determine the optimum algorithm (see Methods). The differences between the top three assemblers (HGAP, Canu and MaSuRCA) were minimal. We reported MaSuRCA as the approach that generally generated the best results, except for the Pf2004 assembly. However, there was only 1 contig of difference between HGAP (20 contigs) and MaSuRCA (21 contigs), and the number of annotated genes and pseudogenes was closer to the reference in the case of the MaSuRCA assembly (5,701 genes and 156 genes) than in the HGAP assembly (5,735 and 205 pseudogenes) (Supplementary Table 1). Table 1 summarizes the information and genome statistics for the assemblies. Despite choosing the best assembler, our results still show that the number of contigs (up to 259) was higher than expected based on the curated Pf3D7 reference (14 chromosomes, 1 Mitochondrion and 1 Apicoplast). A wide range of 5,067-6,210 and 154-449 pseudogenes were also annotated, contrasting with the 5,720 genes and 158 pseudogenes annotated in the Pf3D7 reference, which suggests that further polishing is needed.

### Comparison of methods for the correction of homopolymer tracks

The excessive number of annotated genes and pseudogenes in the *de novo P. falciparum* assemblies (up to 6,210 genes and 449 pseudogenes, Table 1) compared to the 5,720 genes and 158 pseudogenes in the 3D7 reference (Otto, et al., 2018) emphasizes the importance of polishing, as these pseudogenes are probably due to sequencing errors. Homopolymer tracks in long read sequencing technologies are known to give rise to artificial frameshifts, which cause the annotation of truncated gene models and pseudogenes. These occur because long read technologies have issues detecting the correct number of bases in homopolymer tracks or AT repeats. Due to its low GC content (around 19%), tracks of A’s and T’s of over 15 bp in length are frequent in the case of the skewed *P. falciparum* genome. Figure 1 shows an example of a frameshift error in one of the gene models in our *P. falciparum* samples due to the presence of a homopolymer track.

**Figure 1:**
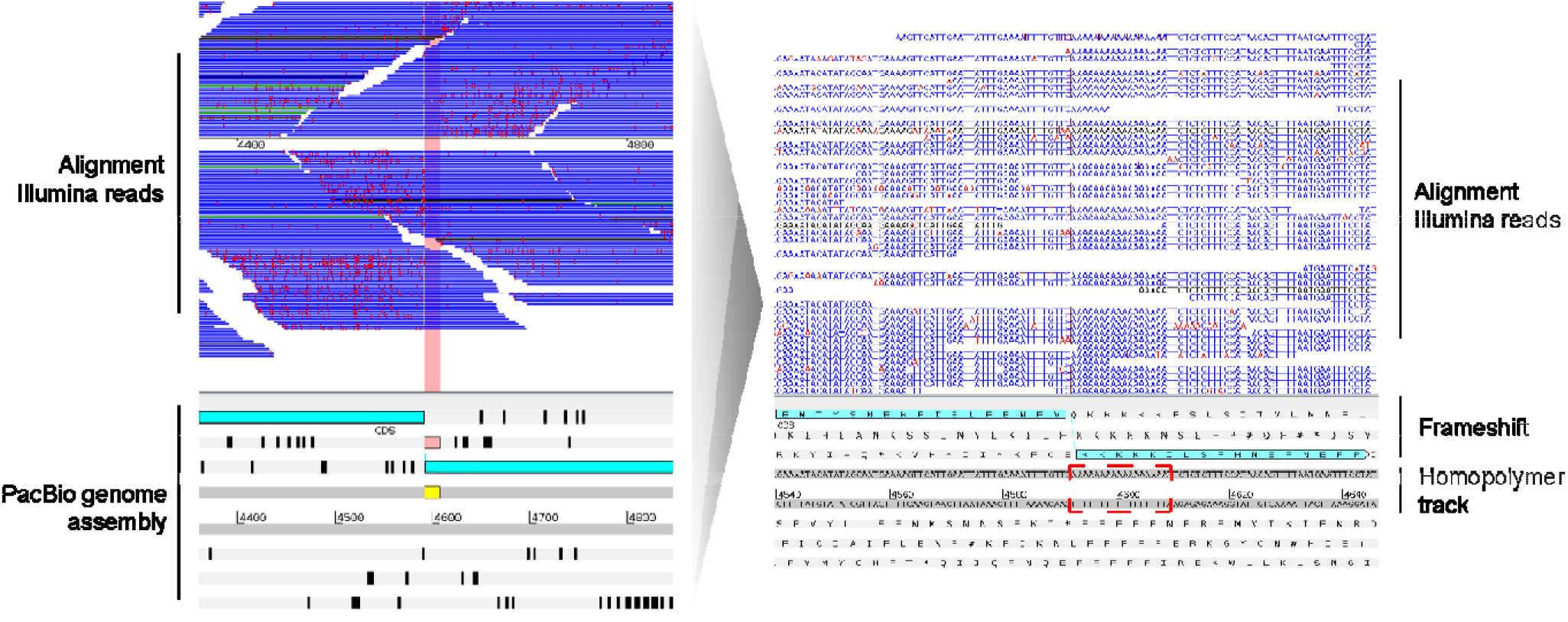
Frameshifts and homopolymer tracks in a long read genome assembly.Artemis visualization of a PacBio genome assembly (bottom panel) and the aligned Illumina short reads (top panel, horizontal blue bars), with reads mapping to the forward strand on top, and to the reverse below. Sequencing errors in the Illumina short reads are marked with vertical light red lines. A homopolymer track of 17 A’s is highlighted in yellow. The quality of the reads drops after the homopolymer, and accordingly it can be seen that reads on the forward strand have just few sequencing errors, but after the homopolymer track the error rate is high. As a homopolymer track is not sequenced correctly, it generates a frameshift and therefore makes a gene model to be wrongly annotated as a pseudogene. In the bottom panel, the two light blue boxes represent exons that due to the indel are split into two. *Ad initio* gene finders could try to build an intron here (loosing exon sequence) or to generate a pseudogene. In the zoom-in visualization (right), the dark red vertical lines in the aligned Illumina short reads point to bases that are missing from the short repetition in the assembly, resulting in the homopolymer track causing the frameshift.

For the ensuing task of correcting homopolymer tracks, we used a comparative approach with the mostly used software, namely iCORN2 and Pilon. We alternatively corrected a long read assembly of the *P. falciparum* 3D7 clone, and compared it to the 3D7 reference (Bohme, et al., 2019) (Table 2 and Supplementary Table 2). When correcting, we also used different sets of Illumina short reads to determine the impact of read length. Both tools were run using alternative aligners in the mapping step (the recommended by the developers and a more sensitive approach with the Bowtie2 aligner), and we evaluated the performance and accuracy of the correction by assessing the number of changes when comparing the corrected sequences to a reference genome (see Methods for details). We argue that comparing the number of pseudogenes in corrected assemblies is a robust method to assess the outcome of tools, as overcorrections are unlikely and the existing functional genes can be confirmed with other approaches, such as protein evidence.

We observed that the iterative approach of iCORN2 when using the Bowtie2 aligner recognizes and corrects more errors than Pilon (Table 2 and Supplementary Table 2). For instance, when correcting with 75 bp Illumina short reads, 531 pseudogenes were annotated in the Pilon-corrected assembly, whereas 164 pseudogenes were annotated in the same assembly after the iterative iCORN2 correction, which is a lower number closer to the 155 pseudogenes from the reference. Overall, around 2-4 times more indels were also corrected with iCORN2 (Supplementary Table 2). Figure 2 shows some examples of the differential correction of frameshifts in the *P. falciparum* 3D7 assembly by Pilon and iCORN2.

**Figure 2:**
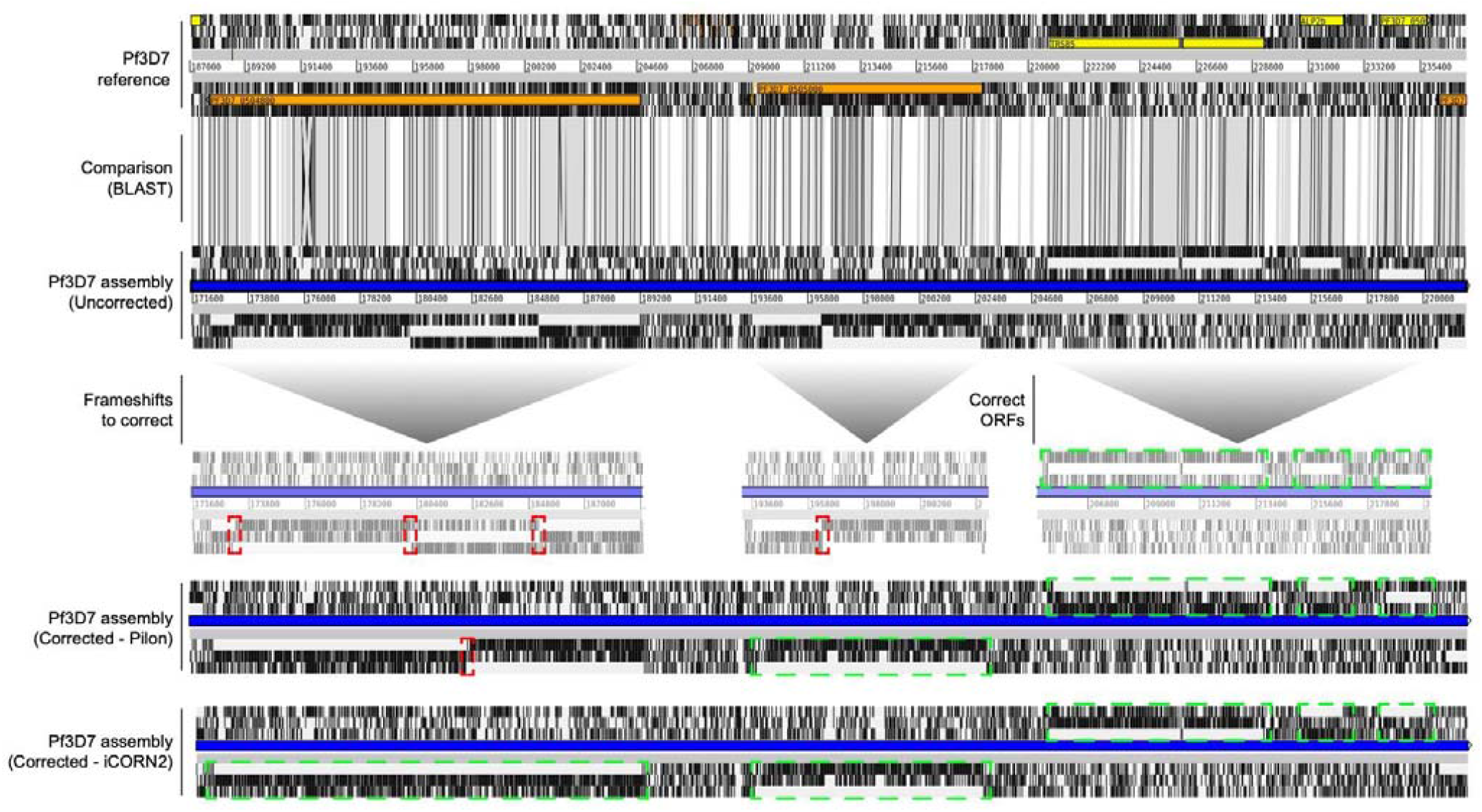
Differential frameshifts correction by Pilon and iCORN2. ACT visualization of a section of the Pf3D7 reference genome, the corresponding section of an uncorrected *P. falciparum* 3D7 PacBio genome assembly, and the Pilon-corrected and iCORN2-corrected sequences. Syntenic regions (BLAST) are indicated in gray bars between the reference and the uncorrected assembly. Annotated genes in the reference are colored. Black vertical lines mark the absence of Open Reading Frames (ORFs). Red squares mark the frameshifts within ORFs in the uncorrected genome sequences. These are differentially processed by Pilon and iCORN2, with multiple iterations of iCORN2 correcting more frameshifts than a single Pilon run. Green squares mark the correct and successively corrected ORFs, which based on the reference could produce proper gene models instead of an excessive and incorrect annotation of pseudogenes.

As expected, the number of corrections also increased with the read length independently of the program, since longer reads can be aligned more accurately and span larger indels. Interestingly, the number of SNPs corrected increased with iCORN2, especially when using longer Illumina short reads (300 bp). Furthermore, 3,919 SNPs and 4,485 indel errors were still uncorrected by iCORN2 compared with the Pf3D7 reference (Table 2 and Supplementary Table 2). As we cannot determine whether this is due to errors in the correction process, the use of different DNA batches, or the presence of errors in the reference, we only used the number of remaining pseudogenes after correction as the best quality metric. Finally, we confirmed that iCORN2 supersedes Pilon by repeating this approach on a *T. brucei* uncorrected assembly. When compared with the *T. brucei* Lister strain 427 2018 reference, around 20% less indels were found in the iCORN2-corrected sequences than in the Pilon-corrected ones, and less pseudogenes were annotated (Supplementary Table 3).

These differences could be due to the iterative implementation of iCORN2. An iterative approach seems to be crucial, and beyond the native implementation of Pilon with a single run, many other studies have successfully corrected genomes using multiple rounds of correction by Pilon (Moser, et al., 2020). Therefore, we also tested the performance of Pilon with the same number of iterations as iCORN2 on the uncorrected *P. falciparum* (100 bp Illumina reads) and the *T. brucei* sequences. We observed that even being ∼5 times slower with default parameters, the annotation of the iCORN2-corrected *P. falciparum* yielded one third less pseudogenes than the Pilon-corrected.

Contrary, for the more fragmented *T. brucei* sequences (Table 3) the Pilon-corrected sequences displayed less pseudogenes, although the differences were very low (∼3%). It should be noted that the high number of pseudogenes in *T. brucei* is due to surface variant genes, which are corrected in an expression cassette to allow antigenic variation. Overall, these observations and the better performance of iCORN2 with *P. falciparum* 3D7 sequences led us to incorporate the iCORN2 software as the default in our pipeline, together with the alternative choice of Pilon.

### Automatic finishing of *de novo* genome assemblies by the ILRA pipeline

As described in more detail in the Methods (Supplementary Figure 1), our ILRA pipeline first cleans short contigs (< than 5,000 bp by default) and contigs that are contained in others by > 90% of their length and with > 99% of identity. Overlaps are then found with a novel method based on merging only if the Illumina coverage is around half of the expected read depth. If a reference sequence exists, the new contigs are also orientated against the reference and renamed. Next, homopolymer errors are iteratively corrected as described above. Finally, the genomes of plasmids and organelle are circularized, the assembly gets decontaminated (discarding contigs representing host or bacterial contamination), and basic statistics are generated, such as the number of potentially complete chromosomes or the completeness of the genomes. ILRA is programmed in Bash and uses internal Bash and Perl scripts.

First, we applied the full ILRA pipeline to the *T. brucei* uncorrected long read assembly used to test iCORN2 and Pilon above (Muller, et al., 2018). The statistics are included in Table 4 and Supplementary Table 1. As expected, the ILRA pipeline successfully corrected *T. brucei* sequences, improving contiguity from 1,232 to 616 contigs (contiguity of the reference genome = 317) and genome size from 65.5 Mbp to 57.8 Mbp (reference genome size = 50.1 Mbp). The number of annotated pseudogenes also decreased by ∼900 (Table 4, Supplementary Table 1).

**Table 4:** Overview of the ILRA improvements step by step in the seven datasets used in this study. Pre-ILRA are the statistics from the assemblies by MaSuRCA, and it is shown the improvement by each step of the pipeline.

| Isolates | *T. brucei* | *L. maculans*<br>*Nz-T4* | *L. biglobosa*<br>*G12-14* | PfCO01<br>RSII | PfCO01<br>Sequel | PfKE07<br>RSII | Pf2004<br>RSII |
| --- | --- | --- | --- | --- | --- | --- | --- |
| **Pre-ILRA** |  |  |  |  |  |  |  |
| #Contigs | 1,232 | 288 | 156 | 30 | 41 | 259 | 21 |
| #Pseudogenes | 5,346 | 540 | 498 | 154 | 163 | 449 | 156 |
| **ILRA correction** |  |  |  |  |  |  |  |
| #Excluded contigs<br><5 kb | 94 | 0 | 0 | 3 | 4 | 3 | 0 |
| #Excluded<br>contained contigs | 254 | 0 | 0 | 0 | 1 | 2 | 0 |
| #Excluded contigs<br>not covered and<br>merged | 33 | 0 | 0 | 0 | 0 | 1 | 0 |
| #Contigs merging<br>and reference<br>ordering | 231 | 186 | 51 | 7 | 12 | 180 | 4 |
| #Excluded contigs<br>contamination | 6 | 1 | 1 | 2 | 7 | 0 | 0 |
| #Contigs (final) | 614 | 101 | 104 | 18 | 17 | 73 | 17 |
| #Pseudogenes<br>(final) | 4,420 | 433 | 420 | 112 | 109 | 223 | 118 |

Next, we demonstrated that ILRA is also applicable to *de novo* genome assemblies from ONT reads. We ran ILRA on two recently published fungal genomes (Dutreux, et al., 2018) and observed improvements in contiguity, annotation of pseudogenes, together with the corrections of thousands of SNPs and indels. While no good reference genomes are available for *Leptosphaeria* spp, contiguity for a *L. maculans* strain Nz-T4 assembly was improved from 288 to 101 contigs and the annotated pseudogenes decreased from 540 to 433, which is closer to the number reported in previous assemblies (Table 4, Supplementary Table 1). Similar improvements were also observed for a *L. biglobosa* G12-14 strain assembly, with 33.3% less contigs and 15.7% less annotated pseudogenes (Table 4, Supplementary Table 1).

To assess the effect of differences in quality and characteristics of the samples and sequences, we generated novel *P. falciparum* assemblies from three isolates with diverse origin: Colombia (PfCO01), Kenya (PfKE07) and Ghana (Pf2004). Table 1 and Table 4 show that the quality of the assemblies obtained using MaSuRCA varied significantly between isolates. The amount of contigs ranged from 21 to 259 and the annotated pseudogenes from 154 to 449. These differences can be attributed to contamination, different median read lengths, different sequencing technologies for PfCO01, or the library preparation protocol, which in the case of PfKE07 was based on whole genome amplification (WGA). We observed that the genomes coming from PacBio RSII reads were finally assembled into a range of 17-73 contigs (median=18, Table 4). These assemblies clearly benefited from the application of ILRA automatic correction, with lower contiguity (previously, range of 21-259 contigs, median=30) and lower number of annotated pseudogenes (Table 4). For example, ILRA allowed for the improved contiguity and the correction of ∼40 gene models incorrectly predicted to be pseudogenes in the Pf2004 sample, which finally assembled into just 17 contigs and had 5,728 genes and 118 pseudogenes annotated. Overall, more than 5,200 genes were annotated in all cases, and contiguity and number of annotated pseudogenes were always improved after ILRA (Table 4, Supplementary Table 1). Moreover, we observed several contigs containing terminal telomeric repeats, which could indicate fully assembled chromosomes, for example 6 out of the 17 contigs of the ILRA-corrected Pf2004 assembly had both telomeres attached (Supplementary Table 1). Despite the general improvements of the ILRA pipeline, there were also some assemblies of lower quality, such as PfKE07. For these sequences, the number of gaps was still high (Supplementary Table 1), and we observed artifacts and mis-assemblies due to the WGA preparation. These errors may be due to the polymerase switching between strands during the amplification step, which results in inverted chimeras generating miss-assemblies that our pipeline was not able to address. Figure 3 displays a schematic case example of an error due to WGA and chimeric reads.

**Figure 3:**
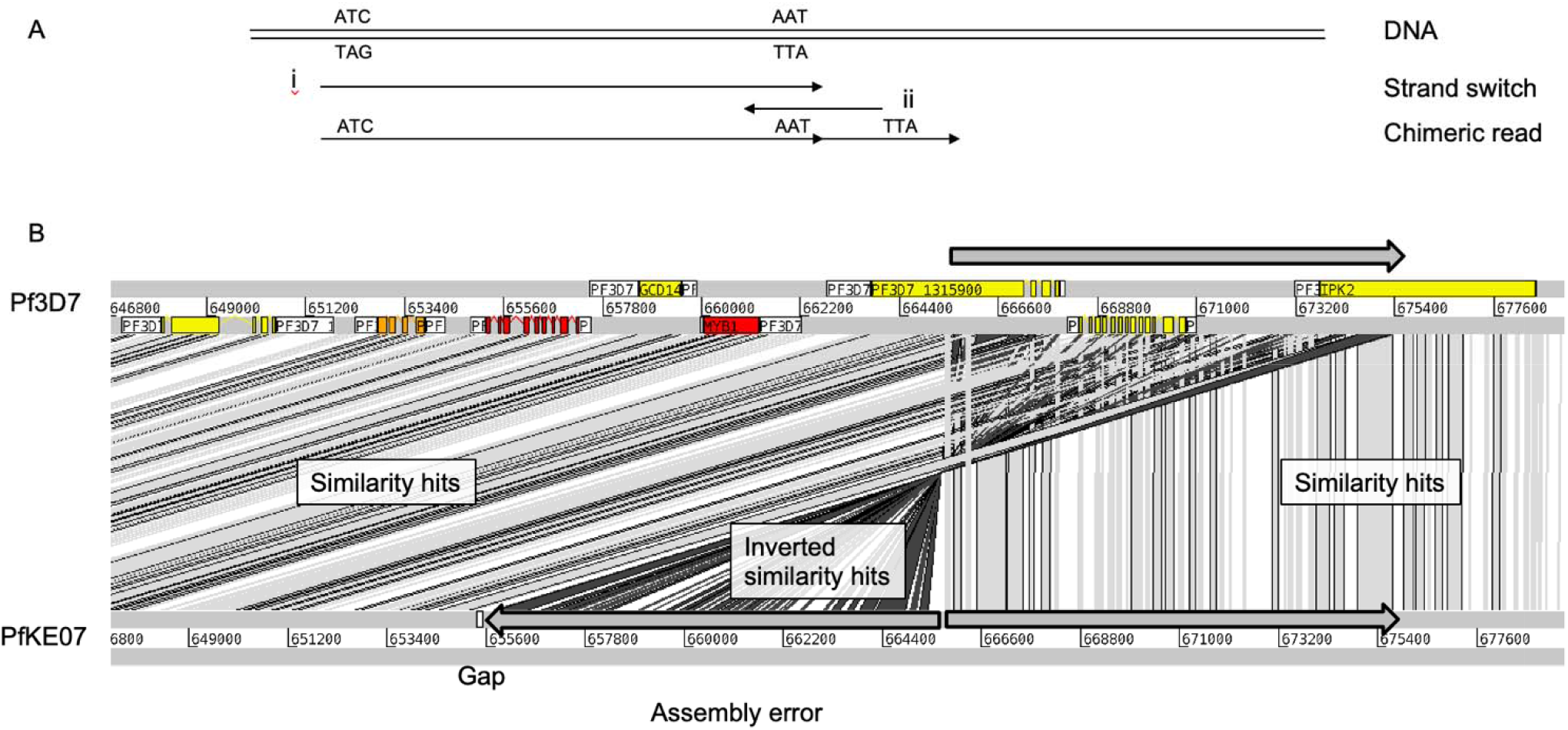
Whole Genome Amplification errors in the PfKE07 assembly. (A)Schematic error of WGA. DNA gets amplified (i), but then the polymerase strand switches and generates the reverse strand (ii). This generates a chimeric read that generates mis-assemblies. (B)These chimeric reads generate assembly errors, as seen in an ACT view. The top part of the reference genome (gray arrow) is duplicated in the WGA amplified genome. The assembly errors generally occur at the contig end, so gaps are generated. Syntenic regions (BLAST similarity hits) when comparing to the reference genome are indicated in gray. Miss-assemblies (Inverted similarity hits) are indicated in black.

Furthermore, to compare the ILRA performance on different types of sequencing reads, we sequenced the library from the *P. falciparum* Colombian sample (PfCO01) using the PacBio Sequel chemistry, in addition to RSII. Of note, this Sequel chemistry was from the first batch of kits, and since then it has been continuously improved by Pacific Bioscience. The initial assembly with the PacBio RSII reads was of considerably higher quality, with approximately 25% more contigs compared to Sequel. After correction by ILRA, the sequences coming from Sequel reads assembled into 17 contigs and similarly, the assembly from RSII reads was composed of 18 contigs (Table 4, Supplementary Table 1). The same library was sequenced with both chemistries, but the mean read length was longer in the RSII (9,413 versus 7,668), while the read depth was higher in the Sequel run (198 versus 168, Supplementary Table 1). Another improvement by ILRA in this case was the identification and removal of multiple contigs identified as *Mycoplasma arginini* contamination (Drexler and Uphoff, 2002), which also caused an excessive number of annotated genes and pseudogenes in both PfC01 assemblies (Table 4).

Finally, we observed that choosing iCORN2 or Pilon within the full ILRA pipeline led to variable but similar numbers of annotated pseudogenes. For the PfCO01 from the Sequel run and the Pf2004 assemblies, the annotation differed in only 1 pseudogene, while the PfCO01 from the RSII run and the PfKE07 assemblies differed in 8 pseudogenes (6.7% less in iCORN2 correction) and 12 pseudogenes (5.1% less in iCORN2 correction), respectively (Supplementary Table S1).

For the last performance test, we also assessed whether ILRA was limited by the size of the genome and/or the length of the sequences. We simulated larger genomes by taking chunks of the human genome with different numbers of contigs and various sizes: ∼100 Mbp, ∼150 Mbp, ∼300 Mbp, 500 Mbp and 1 Gbp (see Methods). We report that ILRA scaled well and successfully processed larger sequences, requiring larger running times and slightly more memory usage (Supplementary Table 4).

### Comparison between the ILRA pipeline and similar tools for finishing genomes

For two of the *P. falciparum* novel genomes (PfCO01 from the RSII run and Pf2004) we also compared ILRA with some existing alternative software for the assembly and correction of sequences, called ARAMIS, Assemblosis and MpGAP (Table 5 and Supplementary Table 1). In the case of PfCO01, we reported an increase on the number of annotated pseudogenes of 36, 34, and 15 for the ARAMIS-, Assemblosis- and MpGAP-processed sequences, which correspond to increases of 32.1%, 30.4% and 13.4% pseudogenes when compared with the ILRA-corrected assembly, respectively (Table 5, Supplementary Table 1). The contiguity post-ILRA was also better (18 contigs vs. 30, 34 and 35 after ARAMIS, Assemblosis and MpGAP, respectively). The number of annotated genes was ∼5,500 for the ILRA-corrected assembly and ∼5,800 after Assemblosis, while it reached 6,200 and 6,300 after ARAMIS and MpGAP. As to Pf2004, when compared with ILRA, the assemblies generated by ARAMIS, Assemblosis and MpGAP displayed very similar numbers of annotated genes (∼5,700), but more contigs (17 contigs in ILRA vs. 21, 22 and 29, respectively), and they contained 20.3%, 40.7% and 9.3% more annotated pseudogenes, respectively (Table 5, Supplementary Table 1).

**Table 5:** Overview of the *P. falciparum* sequences annotated and alternatively assembled and corrected by different tools and pipelines, namely Assemblosis, MaSuRCA+ARAMIS, MpGAP, and MaSuRCA+ILRA.

| Isolates / Tools | PfCO01 RSII |  |  |  | Pf2004 |  |  |  |
| --- | --- | --- | --- | --- | --- | --- | --- | --- |
|  | Assemblosis | ARAMIS | MpGAP | ILRA | Assemblosis | ARAMIS | MpGAP | ILRA |
| **Number of contigs** | 34 | 30 | 35 | 18 | 22 | 21 | 29 | 17 |
| **Size (Mb)** | 23.05 | 23.36 | 23.57 | 22.64 | 23.43 | 23.30 | 23.32 | 23.30 |
| **Number of genes** | 5,763 | 6,159 | 6,298 | 5,521 | 5,743 | 5,718 | 5,707 | 5,723 |
| **Number of** | 146 | 148 | 127 | 112 | 166 | 142 | 129 | 119 |
| pseudogenes |  |  |  |  |  |  |  |  |

## DISCUSSION

With the advent of long reads technologies (Editorial, 2023), the rapid drop of costs and the development of associated algorithms for the analysis, genome sequencing has become more popular and accessible for many laboratories worldwide. Here we demonstrate that automatic finishing of the assemblies is essential to improve the quality of the final genomes. Finishing includes processing such as the removal of small contigs, circularization of organelles with DNA (e.g. mitochondria and apicoplast), renaming of sequences, and addressing errors due to homopolymer tracks and the resultant frameshifts (i.e. polishing). These steps are likely challenging to laboratories without deep bioinformatics knowledge. Here, we develop ILRA, a user-friendly pipeline that combines existing and new tools performing these post-sequencing steps in a completely integrated way, providing fully corrected and ready-to-use genome sequences. We compare its application and performance on existing assemblies from different organisms, as well as novel genome assemblies obtained using various sequencing methods. We show that in all cases ILRA largely improved the sequencing outcome, despite the lower quality of some of the original assemblies.

We applied ILRA to multiple novel and published genomes, including challenging parasites and fungi, both from PacBio and ONT technologies. Our results show that MaSuRCA and HGAP are the best assemblers for *Plasmodium* sequences. MaSuRCA is based on a hybrid approach using both long reads and short reads, while HGAP is a long reads-based assembler officially supported by PacBio as part of SMRT Link, being used to successfully generate assemblies from several *P. falciparum* isolates in a previous study (Otto, et al., 2018). Different samples and tools were of great influence and the assemblies largely differed, but beyond the choice of the assembler software, our comparisons explicitly show that post-assembly polishing steps are often needed. We report that IRLA always improves the contiguity of the assemblies and genome sizes, as well as reducing the number of wrongly assigned pseudogenes. We advocate that it represents an excellent tool for the community to automatically improve long read genome assemblies.

A few other pipelines to automatically finish genomes are already available. Our results show that ILRA outperformed all of them, namely ARAMIS, Assemblosis and MpGAP (Table 5). In the case of Aramis, this is likely due to the fact that it does not manipulate the assembled sequences, such as processing or decontaminating contigs, and only performs polishing using an implementation of Pilon. Assemblosis is an assembly pipeline that uses Canu, and then removes contamination using Centrifuge and polishes using long reads by Arrow and short reads by Pilon. The approach by MpGAP, while implementing different alternatives, is still limited and lacks a decontamination step, which we show may be required for some sequences. Regarding polishing, MpGAP applies an iterative implementation of Pilon via Unicycler to the results of various assembler software. The number of iterations performed is variable, and it increases until it is automatically determined that the best results have been achieved. However, the assembler software included in MpGAP are using only long reads, and, similar to our observation with Canu, HGAP, and Wtdbg2 above, are not necessarily performing the best for challenging organisms such as *Plasmodium*. Indeed, our MpGAP runs with several of the supported assemblers, including Canu, Flye, Raven, Unicycler or Wtdbg2, required dozens of Pilon iterations (up to ∼70), with the better results corresponding to Flye after up to 40 Pilon iterations. However, the numbers of annotated genes and pseudogenes were always improved by our recommendation of MaSuRCA + ILRA with 3 iCORN2 iterations. This highlights the limits of polishing and adds evidence to our observation above that a careful selection of the assembler software is required and more important than polishing, with hybrid approaches such as MaSuRCA performing better for *P. falciparum* sequences. Additionally, ILRA includes unique features, such as the decontamination step (also present in Assemblosis), the reordering of contigs based on a reference, the removal of sequences not covered by Illumina short reads, the analysis of telomeres sequences, or the automatic formatting and filtering following NCBI’s Foreign Contamination Screen, which are not present in any other tool.

We further explored the polishing step to correct sequencing errors with Illumina reads. It is crucial when correcting long read assemblies from PacBio and ONT reads, as these errors may greatly hamper downstream functional annotation. Comparing the existing software for this purpose, we show in *P. falciparum* and *T. brucei* that iCORN2 matches or supersedes Pilon. Even at the cost of larger running time, it seems an iterative approach should always be implemented when improving long reads genome assemblies, in order to make use of the full capabilities of short reads. Thus, the sequences can be iteratively corrected until enough errors are fixed, and Illumina short reads are aligning better. We tested this for Pilon (Supplementary Table 1 and 2), as done by others (Koren, et al., 2017; Naquin, et al., 2018; Tan, et al., 2018). We give the option to the user, who can choose which correction algorithm to use within ILRA. The fact that there is not really a ground truth for a perfect genome highlights the outstanding dilemma that for some reference genomes it is difficult to determine the final consensus sequence. Here we report the correction of pseudogenes as a valid metric, as we have additional evidence that the sequences are correct.

Apart from benefiting from the polishing by ILRA, our *P. falciparum* sequences also illustrate the importance of the initial selection of the sequencing technology. For instance, we observed differences in the PfCO01 assembly from reads by PacBio Sequel or PacBio RSII technologies. Also, despite achieving great improvement and better quality post-ILRA, the PfKE07 assembly was always too fragmented due to the WGA library preparation including artifacts. WGA has been traditionally required to increase the amount of DNA for sequencing in challenging studies. Notably, PacBio recently generated new methods to obtain long reads from 100 ng of DNA (Kingan, et al., 2019), which could render WGA methods obsolete if both high molecular weight DNA and the required expertise are available. However, this is not always possible due to the frequent difficulties to obtain adequate samples when DNA is limited or contaminated by host material, hence automatic polishing tools like ILRA are going to be required. Our four *de novo P. falciparum* assemblies from field isolates, generated by MaSuRCA and automatically corrected by ILRA in this study, are now publicly available. In particular, the isolate from Colombia is the first reference genome recently cultured from South America. We also successfully tested the ILRA pipeline on chunks of human genome up to 1Gbp.

## CONCLUSIONS

Long read technologies enable generation of almost perfect *de novo* genome assemblies from any organism. However, consensus sequences still need polishing and correction of homopolymer errors. Further, in many cases it is not possible to assemble reads at the chromosome level due to limited amount of DNA, low DNA quality or host contamination. In all these cases, the ILRA pipeline is an easy-to-use and accessible tool for any laboratory without deep bioinformatics knowledge, which automatically performs several polishing steps and successfully improves assemblies, making them more continuous and decreasing the number of wrongly assigned pseudogenes.

## Data availability

ILRA can be downloaded from GitHub: https://github.com/ThomasDOtto/ILRA. The code and the *de novo P. falciparum* assemblies corrected in this study are also available in Zenodo - 10.5281/zenodo.7516750. Accession numbers of the published data are, *T. brucei*: ERR1795268/SRR5466319, *L. maculans* NzT4: PRJEB24469, *L. maculans* G12-14: PRJEB24467, and *P. falciparum* reads: ERS037841, ERS001369 and ERS557779, for the 75bp, 100bp and 300 bp, respectively. Novel data can also be found on online databases (accession numbers for long reads, short reads, and the *de novo* assemblies in this study, respectively); Colombian (PfCO01, RSII long reads): ERS2460039, ERS1746432, GCA_019802425.1, Colombian (PfCO01, Sequel long reads): ERS2460039, ERS1746432, GCA_019802405.1, Kenyan (PfKE07): ERS2026796, ERS166385, GCA_019802515.1, and Ghanaian (Pf2004): ERS1412916, ERS1306150, GCA_019802415.1. The corrected assemblies and annotation can also be found on the IRLA GitHub page.

## Keypoints

- The genome sequences resulting from different library preparation approaches, sequencing techniques, and assembler software are variable, and still require polishing and extensive finishing.
- Current software available to fix long-reads sequencing- and homopolymer-related errors may be limited and not fully correct the sequences from different species.
- ILRA is a novel easy-to-use pipeline combining novel and existing tools to automatically performs post-assembly steps and successfully improves sequences.
- The pipeline involves multiple steps and successfully corrects assemblies from different software and species, outperforming existing solutions.

## Acknowledgments

We would like to thank the patients who contributed samples and the health workers who assisted with the sample collections. We would also like to thank staff from the Illumina Bespoke Sequencing Team at the Wellcome Sanger Institute for their contribution. This paper is published with permission from the Director of Kenya Medical Research Institute (KEMRI).

## Figures

**Supplementary Figure 1:**
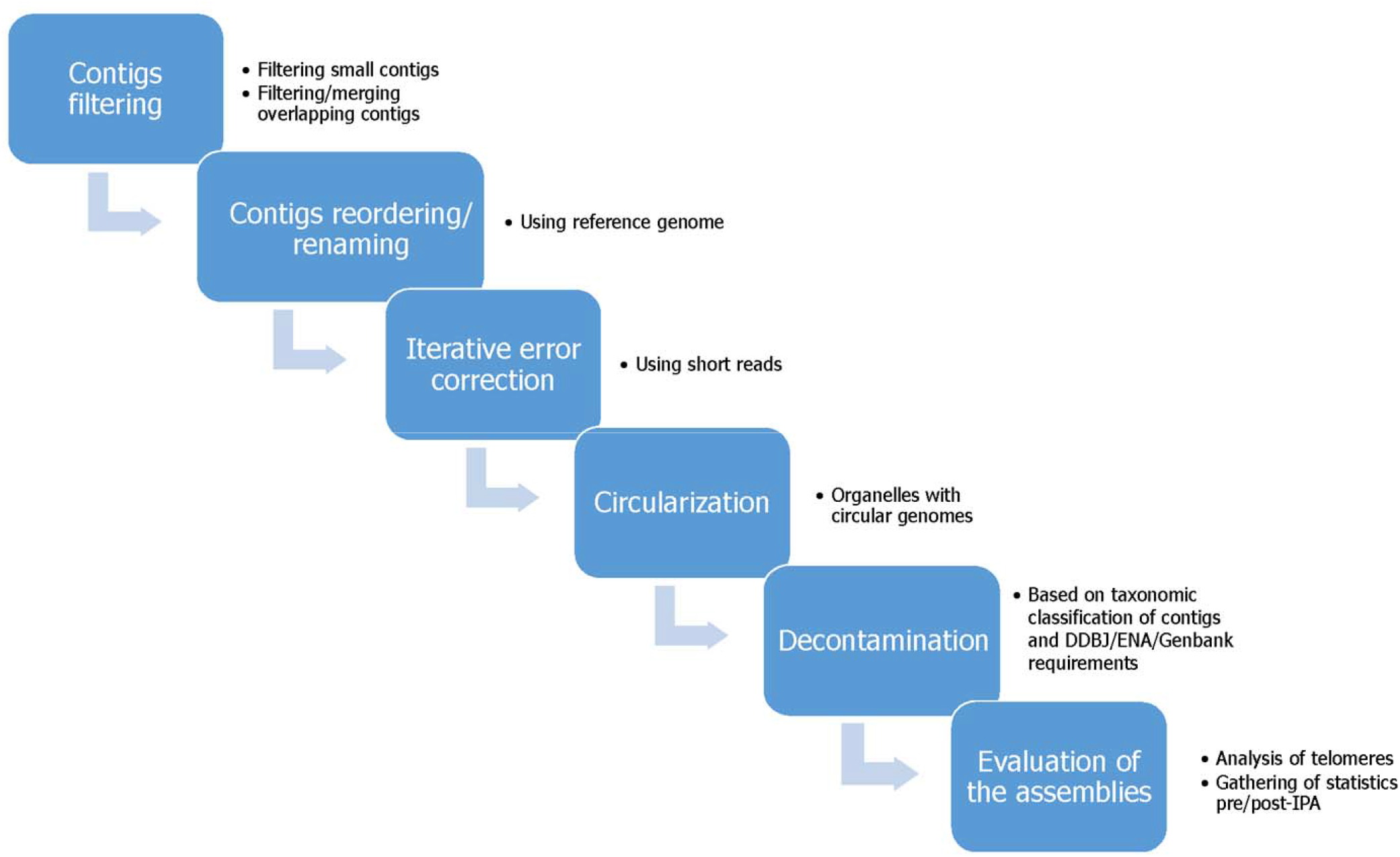
ILRA pipeline workflow.Descriptive flowchart summarizing the steps implemented in the ILRA pipeline: contigs filtering, reordering and renaming, iterative error correction, circularization, decontamination and evaluation of the genome assemblies.

## SUPPLEMENTARY METHODS

### Ethical approval

For all samples written informed consent was obtained directly from adult subjects and from parents or other legal guardians of all participating children, and additional assent was received from children themselves if they were 10 years of age or older. The use of the Kenya parasite isolate (PfKE07) was approved under SSC 1131 that was reviewed and approved by the KEMRI Ethics Review Committee and University of Oxford Tropical Research Ethics Committee. The Pf2004 sample is a culture-adapted strain from Ghana described in (Elliott, et al., 2005). The Colombian sample was an isolate adapted to continuous culture and the keep it in liquid Nitrogen when reached parasitemia >3%. Vials were thawed and DNA extracted using the QIAamp DNA Blood Midi Kit - QIAGEN.

### Sample preparation and sequencing

The samples used in this study are listed in Table 1. The four new isolates were all cultures maintained under standard conditions (Trager and Jensen, 2005). Sequencing by Illumina and Pacific Biosciences (PacBio) SMRT technologies was performed as previously described in (Otto, et al., 2018).

